# Rapid adaptation and remote delivery of undergraduate research training during the COVID 19 Pandemic

**DOI:** 10.1101/2021.02.24.432694

**Authors:** Joanna Yang Yowler, Kit Knier, Zachary WareJoncas, Shawna L. Ehlers, Stephen C. Ekker, Fabiola Guasp Reyes, Bruce F. Horazdovsky, Glenda Mueller, Adriana Morales Gomez, Amit Sood, Caroline R. Sussman, Linda M. Scholl, Karen M. Weavers, Chris Pierret

## Abstract

COVID-19 continues to alter daily life around the globe. Education is particularly affected by shifts to distance learning. This change has poignant effects on all aspects of academic life, including the consequence of increased mental stress reported specifically for students. COVID-19 cancellations of many summer fellowships and internships for undergraduates across the country increased students’ uncertainty about their educational opportunities and careers. When the pandemic necessitated elimination of on-campus programming at Mayo Clinic, a new program was developed for remote delivery. Summer Foundations in Research (SFIR) was drafted around 4 aims: 1) support the academic trajectory gap in research science created by COVID-19; 2) build sustainable scientific relationships with mentors, peers, and the community; 3) create opportunities for participants to share and address concerns with their own experiences in the pandemic; and 4) provide support for individual wellbeing. SFIR included research training, but also training in communication through generative Dialogue and resilience through Amit Sood’s SMART program. 170 participants were followed for outcomes in these spaces. Knowledge of and interest in careers involving biomedical research rose significantly following SFIR. Participants’ mean confidence levels in 12 Key areas of research rose between 0.08 to 1.32 points on a 7-point scale. The strongest gains in mean confidence levels were seen in designing a study and collaborating with others. SFIR participants demonstrated gains in perceived happiness, and measured resilience and a reduction in stress. Participants’ qualitative responses indicated exceptionally positive mentor relationships and specific benefit of both the SMART program and Dialogue.

## Introduction

The emergence and spread of COVID-19 continues to alter daily life around the globe. Education is particularly affected by shifts to distance learning and the retooling of school campuses. This change has poignant effects on all aspects of academic life, including the consequence of increased mental stress reported both in the general population (*1–6*) and specifically for students (*7–10*). Effects on college students have been widely covered in news media (*11–13*). The Healthy Minds Study detailed increasing financial stress among college students, as well as worsening depression, academic impairment attributable to mental health, and rising concerns about the future (*14*).

COVID-19 cancellations of many summer fellowships and internships for undergraduates across the country increased students’ uncertainty about their educational opportunities and careers. These programs serve as critical experiential learning tools for many professional development pathways (*15, 16*). Science, technology, engineering, and mathematics (STEM) summer research fellowships vary by institution, but each plays a key role in student exposure to STEM fields, fosters future opportunities for student professional growth, and boosts recruitment of students to higher learning programs like medical and graduate school. Coupled with a wider education system attempting to adapt to the pandemic, the loss of summer programs heightens the existing vulnerability of pathways to STEM careers and demonstrates the need for innovative programming to ensure the continuity of pathways to postgraduate STEM training.

For the past 30 years, Mayo Clinic has offered 10-week summer undergraduate research programs for students interested in biomedical research training. Typically, students a) participate in mentored laboratory research and career development workshops, b) network with peers, laboratory personnel, and faculty, and c) develop research communication skills. When the pandemic necessitated elimination of on-campus programming, a new program was developed for remote delivery. This 4-week experience, Summer Foundations in Research (SFIR), provided the same core academic pillars of a hands-on fellowship while also addressing documented mental health concerns in the participating undergraduate population (*10*). Program evaluation and an embedded clinical study were included to evaluate achievement of these goals.

### Program objectives

Objectives for the program were drafted around 4 aims: 1) support the academic trajectory gap in research science created by COVID-19; 2) build sustainable scientific relationships with mentors, peers, and the community; 3) create opportunities for participants to share and address concerns with their own experiences in the pandemic; and 4) provide support for individual wellbeing given widespread student mental health concerns both preceding and in relation to COVID-19 (*10*). Participants consisted of a self-selected subset of students previously accepted for in-person summer undergraduate research programs across Mayo Clinic’s three campuses. Original selection criteria for in-person programming required students to have completed at least one year at a US college or university, have a grade point average of at least 3.0 on a 4.0 scale, and have demonstrated interest in biomedical research. Applications were submitted by March 1, 2020, and selection was based on academic experience, research experience, a personal statement, and letters of recommendation. Of 270 eligible students from the initially selected pool, 170 students opted to participate.

### Curriculum development and adaptation for remote delivery

A team of education leaders created the SFIR curriculum by adaptation of existing components and de novo curricular design. The resulting curriculum consisted of four components: introduction to experimental design, Dialogue methodology for communicating science, scientific mentoring, and Stress-Management and Resiliency Training (SMART) (*17, 18*). This curriculum was then adapted for remote delivery via a longstanding Mayo Clinic collaboration with the Integrated Science Education Outreach (InSciEd Out) Foundation (*19, 20*). SFIR was delivered through a combination of synchronous interactions (scientific presentations, small group discussions and one-on-one mentoring) and self-paced asynchronous online modules. The syllabus for the program is included in Supplementary Material 1.

The capstone product of the SFIR experience was a presentation at a virtual poster session. Participants summarized their work with lab immersion mentors and their choice of other personally impactful elements from the four-week program. Posters were presented asynchronously using a five-minute screen capture recording with information uploaded to a shared video database (flipgrid.com/mcsfir2020). Invitations to attend this virtual poster session were emailed broadly at the institutional level and also sent out to families, friends, and mentors of program participants. Poster viewers could also connect directly to Q&A rooms hosted by the presenters for synchronous discussion.

### Outcomes evaluation

The evaluation plan for SFIR was designed to gather critical quantitative and qualitative feedback from participants about the quality and value of each of the program’s components. Key educational outcomes tracked in program evaluation included pre-post changes in career understanding, career interest, and confidence in the development of research skills. Career metrics were evaluated via de novo questions, while research skills confidence was assessed via an adapted subset of 12 items from the Clinical Research Appraisal Inventory(*21*). Further details on the survey methodology and analysis are available in Supplementary Material 2.

The embedded clinical study of wellbeing utilized three questionnaires administered pre-post programming to assess effects upon mental resilience (Brief Resilience Scale (*22*)), stress (Perceived Stress Scale (*23*)), and life satisfaction (Satisfaction with Life Scale (*24*)). These questionnaires and the domains they measure were selected due to their validated psychometric properties (*22–24*), established relevance to SMART training (*17, 25, 26*), and their associations with academic, career, and personal success (*27, 28*). An expanded methods section detailing number of items, validity testing, scoring, cut-off scores, and statistical analyses can be found in the Supplementary Material 2.

## Results

### Educational Outcomes

Participants’ knowledge of and interest in careers involving biomedical research rose significantly following SFIR. The proportion of participants indicating they were “very” or “extremely” knowledgeable about such careers jumped from 16% at baseline to 61% at program end. Inclusion of participants “moderately” knowledgeable of careers in biomedical research pushed this statistic to 99% post- SFIR. Parallel to this trend, the proportion of participants indicating high levels of interest in pursuing biomedical research careers (5 or 6 on a 0 to 6 scale) increased substantially over the course of the program, starting at 33% and ending at 73%. Finally, at program end, 85% of participants indicated they were considering applying to a Mayo Clinic education program; only 4% responded that they were not considering applying, with the remaining 11% reporting uncertain application plans.

Most notable were the gains participants made in confidence across the 12 key research skills measured. Across all skills, participants’ mean confidence levels rose between 0.08 to 1.32 points on a 7-point scale (Figure 1). The strongest gains in mean confidence levels were seen in designing a study and collaborating with others.

**Figure 1.**
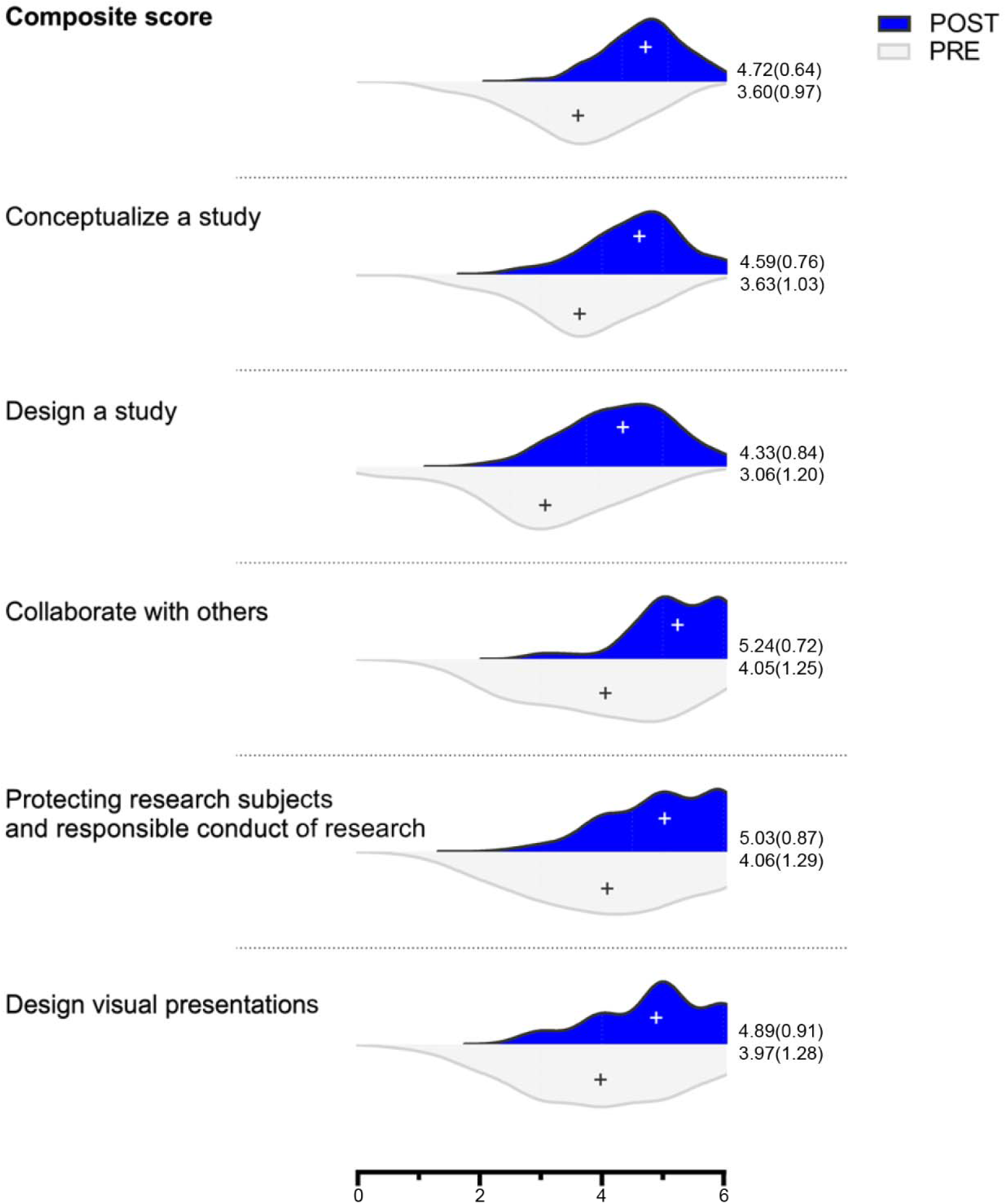
12 items belonging to 1 of 5 categories were selected from the CRAI prior to the beginning of the SFIR program. Items were rated on a scale of 0 to 6, no confidence at all to total confidence. Mean is indicated as “+” and mean (SD) are given to the right of each distribution. The composite score across 12 items had a mean difference (SD_dif_) of 1.11 (0.79) (n=141). Conceptualizing a study (select a suitable topic area, articulate a clear purpose for the research, refine a problem so it can be investigated) 0.97 (0.94) (n=144); design a study (compare major types of studies, choosing an appropriate design to test hypotheses, select appropriate methods of data collection, design the best data analysis strategy) 1.26 (1.17) (n=144); collaborate with others (consult a senior researcher for ideas, participate in generating collaborative research) 1.20 (1.07) (n=143); protect research subjects and responsible conduct of research (discuss ethical issues in research conduct, identify institutional responsibilities in research conduct) 0.97 (1.05) (n=145); and design visual presentations 0.92 (1.12) (n=145).

The post-program survey of participants gathered quantitative and qualitative feedback about program strengths and areas for improvement. Program components that were particularly highly rated included the SMART sessions and mentoring. SMART sessions were rated “quite” or “extremely” worthwhile by 85% of participants. Respectively, 99% percent and 96% of students indicated that their mentors were supportive and showed genuine interest in their research ideas.

### Wellbeing Outcomes

SFIR participants demonstrated gains across all three dimensions of wellbeing. Responses on the Brief Resilience Scale indicate improved resilience after program participation (*M(SD)* pre-3.30 (0.68), post-3.51(0.68), Δ +0.21(0.55), *t*(128) = 4.41, *p* < 0.0001). At the same time, these learners reported decreases in stress on the Perceived Stress Scale (M(SD) pre-19.98(6.89), post-18.06(6.33), Δ −1.91(5.27), *t*(124) = −4.06, *p* < 0.0001). They also recorded increases in life satisfaction, as measured by the Satisfaction with Life Scale (M(SD) pre-24.10(6.03), post-25.43(6.31), Δ +1.33(4.29), *t*(130) = 3.54, *p* = 0.0005). These results correspond to small Cohen’s *d* effect sizes (*29*) in all dimensions (resilience *d* = 0.38; stress *d* = 0.36; life satisfaction *d* = 0.31). Wellbeing trends utilizing established inventory cut-offs are visualized in Figure 2. There is a desirable shift toward normal to high resilience, low to moderate stress, and general to extreme satisfaction with life.

**Figure 2.**
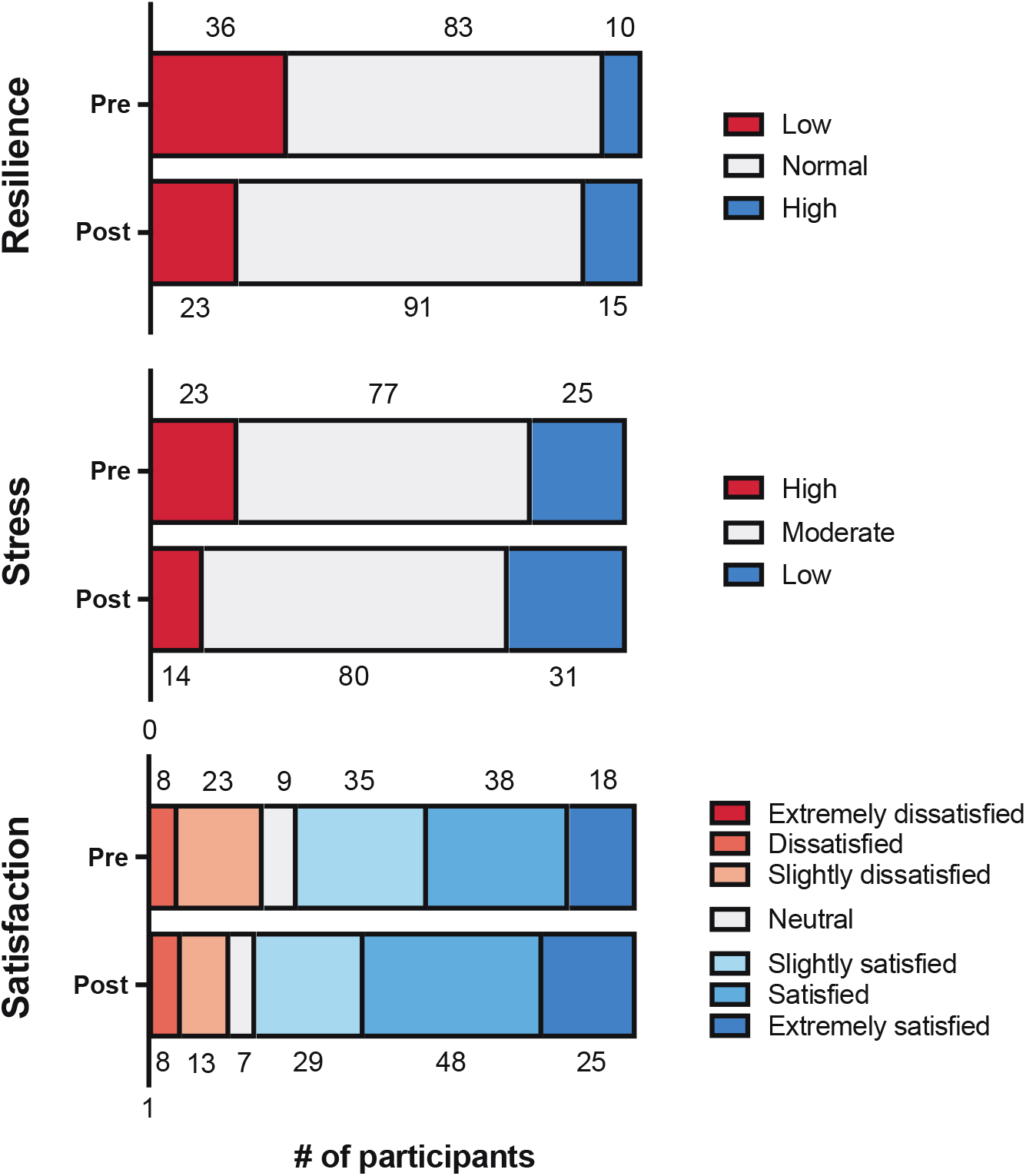
Distribution of participant wellbeing responses pre-post SFIR programming. Score cut-offs for categorization as follows: Brief Resilience Scale N = 129 | Low (1.00–2.99), Normal (3.00–4.30), High (4.31–5.00); Perceived Stress Scale N = 125 | High (27–40), Moderate (14–26), Low (0–13); Satisfaction with Life Scale N = 131 | Extremely Dissatisfied (5–9), Dissatisfied (10– 14), Slightly Dissatisfied (15–19), Neutral (20), Slightly Satisfied (21–25), Satisfied (26–30), Extremely Satisfied (31–35). SFIR students report gains across all three wellbeing categories pre-post programming.

Qualitative assessment of participant feedback found overwhelmingly positive reception of SMART—the programmatic component most explicitly tied to wellbeing. In addition to being a highlight of the program for many participants, SMART’s mindfulness training was interpreted to be important for personal and professional development. Representative comments include:

> *“Mindfulness was incredibly useful because of how it gave me a different perspective on how to address stress and issues in my life”*
>
> and
>
> *“Dr. Sood’s mindfulness sessions were a highlight of the program for me. He gave really concrete and valuable advice for improving relationship(s) and [how to] have a positive, well adjusted mindset. All of these are highly valuable in a scientific career.”*

These comments reinforce results from the wellbeing surveys and indicate the effectiveness of integrating stress-management training into STEM education programming.

## Discussion

Positive educational outcomes for SFIR participants in measurements of career understanding, career interest, and research skills confidence reveal that many goals of research training can be meaningfully addressed in a digital setting. Prior to the delivery of SFIR, Mayo Clinic faculty voiced concerns that the value of summer undergraduate research would be lost without in-person interactions. Major concerns included how participants would develop laboratory skills without setting foot in a laboratory and how mentors would be able to form meaningful connections with young scientists without face-to-face apprenticeships. The results presented herein provide evidence that such fears can be allayed with intentionally designed programming. At the completion of SFIR, participants showed confidence growth in core research skills across all measured domains of study conceptualization, study design, research collaboration, responsible conduct of research, and data presentation. Mentor/mentee relationships flourished in a digital setting. Participants were actively engaged in small peer groups that helped them feel connected to each other and to the program faculty and facilitators. These factors likely contributed to the observed increases in career knowledge and interest.

Improvements in wellbeing metrics of resilience, stress, and satisfaction with life accompanied the above educational gains. This is noteworthy because equipping the next generation of medical and graduate students with tools to decrease stress and improve resilience—two major features of burnout—is of great interest to the academic community (*30–32*). Moreover, a greater satisfaction with life among students is paramount to education as a whole and is a cornerstone for cultivating scientific excellence in a wellness environment (*33*). Although the effect sizes of wellbeing gains were modest, they are like previous interventions that deployed the SMART program (*17, 27*). It is important to note that SMART was a mandatory component of SFIR, which has been shown to restrict effect sizes when compared to opt-in studies (*25*).

To increase confidence in the observed wellbeing results given the turbulent social and political climate of 2020, an external control group of previous Mayo Clinic summer undergraduate students was recruited. Selected students were current sophomores or juniors in the 2020–21 school year and did not receive SMART during their time at Mayo—making them an ideal control for the 2020 SFIR cohort. In comparison to this external control (Supplementary Material 3), SFIR students had statistically significant (Mann-Whitney U test) improvements in resilience (p = 0.03) and decreases in stress (p = 0.03). There was not a significant difference in the gains observed for satisfaction with life (p = 0.81). These results strengthen the assertion that SFIR programming positively impacted student wellbeing even in light of the tumultuous pandemic.

Results from this current study are limited by its participant selection criteria and short-term analysis. Regarding participant selection, SFIR students included a diverse representation (see Supplementary Materials 4) of undergraduates pursuing clinical and translational biomedical research—despite being a selected cohort. In the light of demographic breadth of SFIR participants, study results are likely generalizable to undergraduates engaged in research through US programs. Of interest to education communities at every level is exactly who might struggle and/or benefit the most in a distance-learning environment. Future analyses will consider sociodemographic subgroups of students to address known educational and health disparities. Limitations surrounding length of follow-up and while preliminary data over a three-month period has shown positive outcomes, the sustained impact will be addressed in ongoing longitudinal analyses. This will help elucidate whether there is a need for organizational level maintenance efforts following programs like SFIR. Such data will also provide key insights into equity and inclusion in distance learning for undergraduate students.

Due to the benefits shown in this digital format, SFIR will see continual implementation at Mayo Clinic. The program is being adapted to act as an onboarding experience prior to undergraduate researchers’ arrival for face-to-face mentorship and as a stand-alone offering to increase the reach of Mayo Clinic’s science programming. An abbreviated version preceding summer or school year research aims to enhance the confidence and preparedness of undergraduates for their first experience at a research-focused institution. Furthermore, the program will provide a peer group for support across laboratories, while simultaneously enhancing opportunities for near-peer mentoring by graduate students and postdoctoral fellows.

COVID-19 has challenged learners and educators across the world with its accelerated demand for digital learning. The path of least resistance over the pandemic year has often resulted in cancellation of important educational and professional development programs. While such measures are sometimes unavoidable, a more sustainable approach is to treat COVID-19 as an opportunity for growth—preparing for inevitable future disruptors of the education system. Results from the SFIR case study show that many of the goals of in-person undergraduate biomedical sciences training can be achieved (or even exceeded) in a virtual setting. STEM disciplines thus can embrace adaptation, preserving the integrity of pathways to science, and in doing so, celebrating the spirit of scientific innovation.

## Acknowledgments

The SFIR program was made possible by the William Randolph Hearst Foundation and other benefactors whose generous support of undergraduate research opportunities helps create tomorrow’s physicians and scientists. SFIR relied on the partnership and expertise of the Integrated Science Education Outreach (InSciEd Out) Foundation to convert curriculum for digital delivery. This work was made possible by CTSA Grant Number **UL1 TR002377** from the National Center for Advancing Translational Sciences (NCATS), a component of the National Institutes of Health (NIH). Its contents are solely the responsibility of the authors and do not necessarily represent the official view of NIH.

## References

1. R. Barzilay et al., Resilience, COVID-19-related stress, anxiety and depression during the pandemic in a large population enriched for healthcare providers. Translational Psychiatry 10, (2020).

2. E. A. Holman, R. R. Thompson, D. R. Garfin, R. C. Silver, The unfolding COVID-19 pandemic: A probability-based, nationally representative study of mental health in the United States. Science Advances 6, (2020).

3. L. Rozenkrantz, M. H. Bernstein, C. C. Hemond, A paradox of social distancing for SARS-CoV-2: loneliness and heightened immunological risk. Molecular Psychiatry, (2020).

4. N. Salari et al., Prevalence of stress, anxiety, depression among the general population during the COVID-19 pandemic: a systematic review and meta-analysis. Globalization and Health 16, (2020).

5. D. Vatansever, S. Wang, B. J. Sahakian, Covid-19 and promising solutions to combat symptoms of stress, anxiety and depression. Neuropsychopharmacology 46, 217–218 (2021).

6. J. Xiong et al., Impact of COVID-19 pandemic on mental health in the general population: A systematic review. Journal of Affective Disorders 277, 55–64 (2020).

7. A. K. Cohen, L. T. Hoyt, B. Dull, A Descriptive Study of COVID-19-Related Experiences and Perspectives of a National Sample of College Students in Spring 2020. Journal of Adolescent Health 67, 369–375 (2020).

8. J. F. Huckins et al., Mental Health and Behavior of College Students During the Early Phases of the COVID-19 Pandemic: Longitudinal Smartphone and Ecological Momentary Assessment Study. Journal of Medical Internet Research 22, (2020).

9. J. Lee, Mental health effects of school closures during COVID-19 (vol 4, pg 421, 2020). Lancet Child & Adolescent Health 4, E16–E16 (2020).

10. C. Son, S. Hegde, A. Smith, X. Wang, F. Sasangohar, Effects of COVID-19 on College Students’ Mental Health in the United States: Interview Survey Study. Journal of Medical Internet Research 22, (2020).

11. A. W. June, in The Chronicle of Higher Education. (The Crhinicle of Higher Education, Online, 2020).

12. L. Lumpkin, in The Washington Post. (The Washington Post, washingtonpost.com, 2020).

13. C. Woolston, WHEEL OF FORTUNE: UNCERTAIN PROSPECTS FOR POSTDOCS. Nature 588, 181–184 (2020).

14. A. C. H. Association, “The Impact of Covid-19 on College Student Well-Being,” (Univeristy of Michigan, Healthy Minds Network, 2020).

15. C. Perrett, in Business Insider. (Axel Springer, businessinsider.com, 2020).

16. D. Yaffe-Bellany, in The New York Times. (The New York Times, nytimes.com, 2020).

17. S. S. Chesak et al., Stress Management and Resiliency Training for public school teachers and staff: A novel intervention to enhance resilience and positively impact student interactions. Complementary Therapies in Clinical Practice 37, 32–38 (2019).

18. A. Sood, K. Prasad, D. Schroeder, P. Varkey, Stress Management and Resilience Training Among Department of Medicine Faculty: A Pilot Randomized Clinical Trial. Journal of General Internal Medicine 26, 858–861 (2011).

19. C. Pierret et al., Improvement in Student Science Proficiency Through InSciEd Out. Zebrafish 9, 155–168 (2012).

20. J. Yang, T. J. LaBounty, S. C. Ekker, C. Pierret, Students being and becoming scientists: measured success in a novel science education partnership. Palgrave Communications 2, (2016).

21. E. A. Mullikin, L. L. Bakken, N. E. Betz, Assessing Research Self-Efficacy in Physician-Scientists: The Clinical Research APPraisal Inventory. Journal of Career Assessment 15, 367–387 (2007).

22. B. W. Smith et al., The brief resilience scale: assessing the ability to bounce back. International journal of behavioral medicine 15, 194–200 (2008).

23. K. T. Cohen S, Mermelstein R, et al., Measuring stress: A guide for health and social scientists. S. Cohen, R. C. Kessler, L. U. Gordon, Eds., Measuring stress: A guide for health and social scientists. (Oxford University Press, New York, NY, US, 1997), pp. xii, 236–xii, 236.

24. E. Diener, R. A. Emmons, R. J. Larsen, S. Griffin, The Satisfaction With Life Scale. Journal of personality assessment 49, 71–75 (1985).

25. L. N. Dyrbye et al., The Impact of a Required Longitudinal Stress Management and Resilience Training Course for First-Year Medical Students. Journal of General Internal Medicine 32, 1309–1314 (2017).

26. K. Etherton, D. Steele-Johnson, K. Salvano, N. Kovacs, Resilience effects on student performance and well-being: the role of self-efficacy, self-set goals, and anxiety. The Journal of General Psychology, 1–20 (2020).

27. A. Bhagra et al., Stress Management and Resilience Intervention in a Women’s Heart Clinic: A Pilot Study. Journal of Womens Health 28, 1705–1710 (2020).

28. S. J. Wolf, T. M. Lockspeiser, J. Gong, G. Guiton, Identification of foundational non-clinical attributes necessary for successful transition to residency: a modified Delphi study with experienced medical educators. Bmc Medical Education 18, (2018).

29. J. Cohen, Statistical Power Analysis for the Behavioral Sciences. (Academic Press, 1988), pp. 490.

30. L. N. Dyrbye, W. Lipscomb, G. Thibault, Redesigning the Learning Environment to Promote Learner Well-Being and Professional Development. Academic Medicine 95, 674–678 (2020).

31. R. K. Jordan et al., Variation of stress levels, burnout, and resilience throughout the academic year in first-year medical students. Plos One 15, (2020).

32. M. P. S. Mousavi et al., Stress and Mental Health in Graduate School: How Student Empowerment Creates Lasting Change. Journal of Chemical Education 95, 1939–1946 (2018).

33. E. Amdurer, R. E. Boyatzis, A. Saatcioglu, M. L. Smith, S. N. Taylor, Long term impact of emotional, social and cognitive intelligence competencies and GMAT on career and life satisfaction and career success. Frontiers in Psychology 5, (2014).

