## Supplementary Materials for "Rapid adaptation and remote delivery of undergraduate research training during the COVID 19 Pandemic"

### Summer Foundations In Research 2020

**Program Director**
Chris Pierret, Ph.D.

**Program Administrator**
Karen Weavers, M.Ed.

**SURF Oversight**
Bruce Horazdovsky, Ph.D.

**SURF Coordinator**
Glenda Mueller

**UREP Oversight**
Caroline Sussman, Ph.D.

**Program Instructors**

Biomedical Research
Anthony Windebank, M.D.

Study Design
Jennifer St. Sauver, Ph.D.

Statistical Methods
Stacey Winham, Ph.D.

Communicating Research
Stephen Ekker, Ph.D.

Track Plenaries
Michael Barry, Ph.D.
Aaron Johnson, Ph.D.
Pamela McLean, Ph.D.
Isobel Scarisbrick, Ph.D.
Lisa Schimmenti, M.D.
Anthony Windebank, M.D.
Richard Weinshilboum, M.D.

Experimental Design Small Group Facilitators
Jennifer Arroyo, Ph.D

**Program Overview**

Welcome to this brand-new online course: Mayo Clinic's – Summer Foundations in Research. This four week offering includes 1) an introduction to experimental design; 2) training in dialogue methodology for communicating science with our communities; 3) track specific seminar series; 4) SFIR specific seminar series; 5) one-on-one scientific mentorships; 6) connectedness to the Mayo Clinic research community through lab meeting attendance; and 7) Resilience training through Dr. Amit Sood's SMART program. These opportunities will be further supplemented by online Clinical and Translational Science coursework and video lectures.

The program will conclude with a virtual research symposium where students will have an opportunity to present their work with their mentors from the summer. Students will also have the opportunity to network with their peers and Mayo Clinic Scientists.

**Program Objectives**

- Students will be introduced to the cutting-edge science of Mayo Clinic and the Mayo Clinic research community
- Students will develop a well-rounded understanding of experimental design and scientific inquiry.
- Students will complete a scientific experience to be presented at the virtual symposium that will propel them in their scientific trajectory.
- Students will build resiliency through the completion of Dr. Amit Sood's Resilient Option program.
- Students will learn dialogue skills for use in the communication of their science to their community.

**Core Program Components**

Experimental Design 1 – July 20-29
An introduction to the topic of scientific research, methods, analysis, and presentation with each MCGSBS track discussing their research.

Experimental Design 2 – July 30- August 5

All participants will have the opportunity to work directly with Mayo Clinic researchers in small groups with a small group facilitator by

Arjun Athreya, Ph.D.
Roberto Cattaneo, Ph.D.
Doo-Sup Choi, Ph.D.
Martin Fernandez-Zapico, M.D.
Peter Harris, Ph.D.
Kathryn Knoop, Ph.D.
Kun Ling, Ph.D.
Luis Lujan, Ph.D.
Carmen Reynolds, Ph.D.
Laura Rogers, Ph.D.
Michael Romero, Ph.D.
Caroline Sussman, Ph.D.

Resilient Option
Amit Sood, M.D.
Chris Pierret, Ph.D.

Dialogue Small Group Facilitators
Chris Pierret Ph.D.
Catherine Knier
Fabiola Guasp Reyes
Adriana Maria Morales-Gomez
Carmen Silvano
Karen Weavers M.Ed.
Malachi Johnson
Jim Sonju
Allison Krinke
Kristina Johnson
Will Ruffin
Ting Wei, Ph.D.
Martin Fernandez-Zapico, M.D.
Louis El Khoury, Ph.D.
Santiago Restrepo Castillo

**Program Manager**

Zachary WareJoncas


**Program Director Office Hours**
Every Weekday from 3-4pm.
Office Hours Link

track. The facilitators will provide 2-3 journal articles that represent the research within their track. These will be discussion sessions developing research questions and identifying methods to address those questions.

Direct Mentorship (July 20- August 14)
Participants will have a one-on-one session with a mentor and/or their postdoc, PhD student and others in the lab to learn about individual research projects and to participate digitally in ongoing laboratory research. **The mentor will identify activities for the participant but will not be with them all the time. Times are included in the schedule below, but that is only as a place holder and can occur at any time.**

Presentations and Meetings
Throughout the program there will be opportunities to attend presentations at three different scales.

- **SURF Seminar Series** are available to the entire program.
- **Track Talks** are presentations given within the individual research departments to which you have been assigned, these will include seminars, journal clubs and works in progress (WIP) meetings.
- **Lab Meetings** will enable you to experience the day-to-day science of a lab group at Mayo Clinic.
- **TBA Networking Opportunities** to include no-cost conference opportunities among other event forums.

Dialogue (July 21 – August 11)
Generative Dialogue sessions are designed to create conversational norms and safe space to discuss complex issues, experience vulnerability as a group and communicate effectively with diverse stakeholders. Developing this skillset will be an invaluable tool in communicating science with your community in the future.

Resilient Option (July 29 – August 14)
2020 has been a trying year for a wide variety of reasons and even if this year was not an especially poignant example, careers and science and medicine can be among the most stressful occupations. We are including mindfulness training following Dr. Amit Sood's Stress Management and Resiliency Training (SMART) program as a core component of SFIR to provide training on tools that can be used now and throughout your career to combat stress and distraction.

**Key Resources**

Blackboard
Blackboard is our virtual classroom system for this program. You will be given a user account to log-in for the duration of this program. Resources for learning how to use blackboard can be found at the main blackboard website or by contacting the Program Manager. (https://www.blackboard.com/)

FunCaTs (approx. 4.5hrs)
Fundamentals of Clinical and Translational Science (FunCaTS) is a combination of 13 online modules strategically packaged together to enable medical professionals and allied health staff to expand their knowledge of the components of clinical and translational research. These components provide the fundamental knowledge to promote understanding of the process of bringing discoveries to the bedside and then to the population.

Sample Size Matters (approx. 10hrs)
Sample Size Matters is an online program for graduate students and research professionals. Participants learn to apply the knowledge and skills gained to improve rigor, transparency and reproducibility in their own research.

#### Program Schedule

| Day | Schedule | Activity | Description |
|---|---|---|---|
| **Week 1** | CST | | |
| **Mon July 20th** | 10:00am-12:00pm | Welcome to SFIR | ***Introduction of program faculty and MCGSBS leadership* – Stephen Ekker, PhD** |
| | Track Scheduled | Presentations and Meetings | Determined by track |
| | Independent | Pre-Evaluations | Program Evaluation & Wellness Study |
| **Tues July 21st** | 10:00am-11:00am | Experimental Design 1-1 | ***Why Biomedical Research is Important* – Anthony Windebank, MD** |
| | 11:00am-12:00pm | **Dialogue 1 Section A** | **Introduction to Dialogue** |
| | Track Scheduled | Presentations and Meetings | Determined by track |
| | Independent | **FunCaTS** | At own pace |
| **Wed July 22nd** | 10:00am-11:00am | Experimental Design 1-2 | ***Study Design Methods* – Jennifer St. Sauver, PhD** |
| | 11:00am-12:00pm | **Dialogue 1 Section B** | **Introduction to Dialogue** |
| | Track Scheduled | Presentations and Meetings | Determined by track |
| | Independent | **FunCaTS** | At own pace |
| **Thurs July 23rd** | 10:00am-11:00am | **Resilient Option 1** | **Gratitude** |
| | 11:00am-12:00pm | **Experimental Design 1-3** | ***Statistical Methods* – Stacey Winham, PhD** |
| | Track Scheduled | Presentations and Meetings | Determined by track |
| | Independent | **FunCaTS** | At own pace |

| Day | Schedule | Activity | Description |
|---|---|---|---|
| **Fri July 24th** | 10:00am-11:00pm | **Experimental Design 1-4** | **Track Leadership Plenaries**<br>10:00 - VGT - Michael Barry, PhD<br>10:15– MPET-Richard Weinshilboum, MD |
| | 11:00am-12:00pm | **Experimental Design 1-5** | **Track Leadership Plenaries**<br>11:00-NSC - Pamela McLean, PhD<br>11:15-BMB - Lisa Schimmenti, MD |
| | Track Scheduled | Presentations and Meetings | Determined by track |
| | Independent | **FunCaTS** | At own pace |
| **Week 2** | | | |
| **Mon July 27th** | 10:00am-11:00am | **Experimental Design 1-6** | **Track Leadership Plenaries**:<br>10:00-IMM – Aaron Johnson, PhD<br>10:15- nuSURFs – Michael Romero, PhD |
| | 11:00am-12:00pm | **Dialogue 2 Section A** | **COVID and My Science** |
| | Track Scheduled | Presentations and Meetings | Determined by track |
| | Independent | **Sample Size Matters** | |
| **Tues July 28th** | 10:00am-11:00am | **Experimental Design 1-7** | **Track Leadership Plenaries**:<br>10:00-REGS-Isobel Scarisbrick, PhD<br>10:15-CTS-Anthony Windebank, MD |
| | 11:00am-12:00pm | **Dialogue 2 Section B** | **COVID and My Science** |
| | Track Scheduled | Presentations and Meetings | Determined by track |
| | Independent | **Sample Size Matters** | At own pace |
| **Wed July 29th** | 10:00am-11:00am | **Experimental Design 1-8** | ***Communicating Research* – Stephen Ekker, PhD** |
| | 11:00am-12:00pm | **Resilient Option 2** | **Mindful Presence** |
| | Track Scheduled | Presentations and Meetings | Determined by track |
| | Independent | **Sample Size Matters** | At own pace |
| **Thurs July 30th** | 10:00am-11:00am | **Experimental Design 2-1** | **Small Group Experimental Design** |
| | 11:00am-12:00pm | **Resilient Option Followup Section A** | Group Capacity Building |
| | Track Scheduled | Presentations and Meetings | Determined by track |
| | Independent | **Sample Size Matters** | At own pace |
| **Fri July 31st** | 10:00am-11:00am | **Experimental Design 2-2** | Small Group Experimental Design |
| | 11:00am-12:00pm | **Resilient Option Followup Section B** | Group Capacity Building |
| | Track Scheduled | Presentations and Meetings | Determined by track |
| **Week 3** | | | |
| **Mon August 3rd** | 10:00am-11:00am | **Experimental Design 2-3** | Small Group Experimental Design |
| | 11:00am-12:00pm | **Dialogue 3 Section A** | **COVID and My Community** |
| | Track Scheduled | Presentations and Meetings | Determined by track |
| **Tues August 4th** | 10:00am-11:00am | **Experimental Design 2-4** | Small Group Experimental Design |
| | 11:00am-12:00pm | **Dialogue 3 Section B** | **COVID and My Community** |
| | Track Scheduled | Presentations and Meetings | Determined by track |
| | Independent | Activities on shared calendar | |

| Day | Schedule | Activity | Description |
|---|---|---|---|
| **Wed August 5th** | 10:00am-11:00am<br>11:00am-12:00pm<br>Track Scheduled | **Experimental Design 2-5**<br>**Resilient Option 3**<br>Presentations and Meetings | Small Group Experimental Design<br>**Kindness**<br>Determined by track |
| **Thurs August 6th** | 10:00am-11:00am<br>11:00am-12:00pm<br>Track Scheduled | Mentoring 1<br>**Resilient Option Followup Section A**<br>Presentations and Meetings | Time with individual mentors<br>**Group Capacity Building**<br>Determined by track |
| **Fri August 7th** | 10:00am-11:00am<br>11:00am-12:00pm<br>Track Scheduled | Mentoring 2<br>**Resilient Option Followup Section B**<br>Presentations and Meetings | Time with individual mentors<br>**Small Group Capacity Building**<br>Determined by track |
| **Week 4** | | | |
| **Mon August 10th** | 10:00am-11:00am<br>11:00am-12:00pm<br>Track Scheduled | Mentoring 3<br>**Dialogue 4 Section A**<br>Presentations and Meetings | Time with individual mentors<br>**My Community and My Science**<br>Determined by track |
| **Tues August 11th** | 10:00am-11:00am<br>11:00am-12:00pm<br>Track Scheduled | Mentoring 4<br>**Dialogue 4 Section B**<br>Presentations and Meetings | Time with individual mentors<br>**My Community and My Science**<br>Determined by track |
| **Wed August 12th** | 10:00am-11:00am<br>11:00am-12:00pm<br>Track Scheduled | Mentoring 5<br>**Resilient Option 4**<br>Presentations and Meetings | Time with individual mentors<br>**Resilient Mindset**<br>Determined by track |
| **Thurs August 13th** | 10:00am-11:00am<br>11:00am-12:00pm<br>Track Scheduled<br>Independent | Mentoring 6<br>**Resilient Option Followup Sections A/B**<br>Presentations and Meetings<br>Poster/Presentation Prep | Time with individual mentors<br>**Group Capacity Building**<br>Determined by track |
| **Fri August 14th** | 10:00am-10:30am<br>10:30am-12:00pm | Closing Thoughts<br>**Virtual Symposium** | Final messages and poster session details<br>**Presentation of Posters to Peers and Guests** |

#### Program Evaluation Team

| Education | Wellness |
|---|---|
| Linda Scholl Ph.D. | Joanna Yang-Yowler Ph.D. |
| Karen Weavers M.Ed. | Catherine Knier |
| Adriana Maria Morales-Gomez | Fabiola Guasp Reyes |

**Supplemental Materials: Expanded Methods**

#### Program Design & Delivery

##### Overview/Syllabus

In addition to the program objectives detailed in this paper's main text, structural design goals in creation of SFIR programming were: (1) to provide a robust scientific immersion experience for participants in a virtual environment, (2) to fit this program into a four week period to accommodate pandemic-induced school term changes, (3) to limit synchronous components to two hours a day, (4) to provide many opportunities for small group and 1:1 interactions, and (5) to improve participant wellbeing before their return to regular undergraduate programming. The full program syllabus is provided in Supplementary Material 3 In brief, this schedule was designed to deliver one hour of science-focused programming and one hour of wellbeing or communication-focused programming each morning with afternoons reserved for independent work.

##### Experimental Design

Experimental Design 1 and Experimental Design 2 program components were derived from a curriculum developed via long-standing partnership between the Center for Clinical and Translational Science at Mayo Clinic and the University of Puerto Rico Medical School. The goal of these two curricular modules was to provide a robust introduction to the fundamentals of clinical research (Experimental Design 1) and a mentored immersion in the design of a clinical experiment (Experimental Design 2). Both modules were modified for digital delivery in SFIR. Live video lectures were used for program-wide talks, and continuously available online meeting rooms were created for use by small groups. Lecturers for Experimental Design 1 were recruited from faculty associated with the Center for Clinical and Translational Science, and mentors for Experimental Design 2 were recruited from the faculty, post-doctoral fellows, and graduate students at the Mayo Clinic Graduate School of Biomedical Sciences. Small groups for Experimental Design 2 were limited to a minimum of six and a maximum of ten to ensure participants had similar opportunities for direct interaction with their mentors. Participant groupings were determined by the graduate school tracks each participant would have been associated with had the regular summer undergraduate research program proceeded.

##### Scientific Dialogue

The generative Dialogue sessions were designed according to principles described in Pierret et al.'s 2012 publication (*1*). Dialogue question templates (Supplemental Material 4) guided each session and mapped a series of four interconnected Dialogues initially focused on individual experiences of the COVID-19 pandemic before broadening out into an exploration of community experiences. To accommodate the large class size of SFIR, conversational norms were established primarily in the large group lecture with final adaptation performed in each small group. Dialogue facilitators were recruited from both Mayo Clinic and the InSciEd Out Foundation, resulting in a diverse group of faculty, post-doctoral fellows, graduate students, and public school teachers. Participant groups from Experimental Design 2 were used in Dialogue to foster continued interaction with the same group of peers.

##### Stress-Management and Resiliency Training (SMART)

SMART programming (https://www.resilientoption.com/) was provided by the Global Center for Resiliency Training and has been previously delivered in an online format, so no modification to the primary lessons was necessary. To increase buy-in and facilitate training, follow-up 1-hour Q&A sessions were interspersed between lesson deliveries. These Q&A sessions were hosted in half-class sections to provide more opportunities for each participant to ask questions and comment on the lessons.

Direct Mentorship

The direct mentorship component was modeled after the usual in-person summer undergraduate research program, matching students either 1:1 or in small groups with scientific mentors for an immersion experience. Where possible, participants were paired with the labs that would originally have hosted them during the in-person program. If original labs were not available, students were paired up with another student from the same department for a joint mentorship experience. The mentors could determine the specifics of each mentorship experience within the guidelines outlined in Supplementary Material 5.

Supplemental Digital Learning

The Fundamentals of Clinical and Translational Science (FUNCaTS) and Sample Size Matters courses were originally designed to be self-paced digital offerings, so no modification was necessary.

Additional Lecture Offerings

Further seminars and lectures were offered through the duration of the program. These were delivered in a traditional video lecture format over Blackboard Collaborate, and the design of each session was at the discretion of the lecturer.

Capstone Poster Presentation

The capstone poster session was designed to maximize participant and guest interaction through a combination of asynchronous poster viewing and synchronous Q&A sessions. Posters were hosted on Flipgrid, a short-format video recording site produced by Microsoft. Participants were given a poster template with sections dedicated to Experimental Design, Mentorship, and program experience. They were directed to record five-minute screen share videos on Flipgrid to present their posters. Participants and guests were then invited to view posters during a 1.5-hour period. For the subsequent Q&A sessions, participants were randomly assigned into groups of ten with people not from their experimental design group to ensure presentation variety. Each group was then given a Blackboard Collaborate session to jointly host. During the poster session, participants were assigned posters from other presenters in their Q&A room and instructed to prepare questions. Guests were directed to the Q&A rooms for each poster by links placed in the poster descriptions within Flipgrid. Q&A sessions ran for 1.5 hours, and a member of the program delivery team visited each room at least once.

**Outcomes Evaluation**

Education Outcomes

As mentioned in the main text, education outcomes evaluation focused upon capturing changes in career understanding, career interest, and confidence in the development of research skills.

Participants were asked to complete pre-post surveys: the first at the beginning of the program, and the second on the last day of the program. Participants were given time during the orientation to fill out the survey, and completion of both surveys was required in order to receive the program stipend.

The pre-program questionnaire was brief. One item asked participants to rate their level of knowledge about careers involving biomedical research on a 5-point Likert scale. Participants were also asked to rate their level of confidence on 12 key research skills drawn from the Clinical Research Appraisal Inventory (CRAI). Two validated versions of the CRAI instrument are available (*2, 3*), both of which contain items that either are not appropriate for undergraduate at a very early stage in the development (e.g., recruit and screen research project staff) or are specific to clinical research rather than to basic research (e.g., describe ethical concerns with the use of placebos in clinical research). In order to cull these items and focus upon SFIR program targets, two SFIR program leaders reviewed items from the full CRAI instrument (*2*) and selected twelve most relevant to SFIR's targeted goals. The skills were: 1) selecting a suitable topic area for study; 2) articulate a clear purpose for the research; 3) refine a problem so it can be investigated, 4) compare major types of studies (e.g., case controls, cohort/longitudinal studies, clinical trials); 5) choose an appropriate research design that will answer a set of research questions and/or test a set of hypotheses; 6) select methods of data collection appropriate to the study; 7) design the bet data analysis strategy for a study; 8) consult senior researchers for ideas; 9) participate in generating collaborative research ideas; 10) discuss ethical issues involved in conducting research; 11) identify the responsibilities of research institutions in conducting research; 12) design visual presentations (posters, slides, graphs, pictures). The items asked participants to rate their level of confidence performing each skill on a seven-point scale (from no confidence to total confidence. The published CRAI instruments use a 10-point confidence scale; however, the evaluation team decided to use a 7-point scale due, again, to the relatively early stage of research development of the SFIR participants.

The end-of-program survey again asked students to rate their knowledge of and interest in careers involving biomedical research, as well as their confidence on the 12 key research skills. In addition, the survey included a course evaluation section that asked participants to rate how worthwhile each of the program components were (on a 5-point Likert scale) and provide some narrative comments and suggestions for program improvement. Finally, participants were asked about their future educational plans, including whether or not they were considering applying to a Mayo Clinic education program in the future. Their responses were captured in Qualtrics and compiled into a comprehensive spreadsheet that included participant-level demographics (e.g., gender, race/ethnicity, first generation college status, economic/academic disadvantage, disability status, and domestic/international status). These demographic variables will be analyzed further in subsequent papers.

<u>Wellbeing Outcomes</u>

As mentioned in the main text, wellbeing was assessed via a battery of three questionnaires, which covered mental resilience (Brief Resilience Scale), stress (Perceived Stress Scale), and life satisfaction (Satisfaction with Life Scale). Measures were selected to span dimensions of interest, and each is widely used and well-validated in undergraduate and adult populations (see specific references below). The wellbeing study was reviewed and approved by Mayo Clinic's COVID-19 Committee and Education Committee. Subsequently, the Institutional Review Board approved study activities as minimal risk research exempt under 45 CFR 46.101 Category 2. Assent was sought before the start of each wellbeing survey.

The Brief Resilience Scale (BRS) is a six-item assessment of self-reported resilience scored on a five-point Likert scale (*4-6*). It has previously been implemented in evaluations of InSciEd Out (data pending publication) and SMART (*7*). The summative score is the average across all six items (1–5) with higher averages indicating higher resilience. Categorical characterization of average score spans Low (1.00–2.99), Normal (3.00–4.30), and High (4.31–5.00) resilience.

The Perceived Stress Scale (PSS) consists of 10 items that assess perceptions of stress on a five-point Likert scale (*8-11*). It queries thoughts and feelings over the past month, making it an ideal measure for the roughly month-long SFIR program. It has also previously been used in evaluations of SMART (*12, 13*). The summative score is the total of all ten items (0–40) with lower scores indicating lower stress. This score is divided into categories of: High (27–40), Moderate (14–26), or Low (0–13) stress.

The Satisfaction with Life Scale (SWLS) is a five-item self-reported measure of an individual's current happiness scored on a seven-point Likert scale (*14-16*). It has been validated and performs similarly whether delivered in pencil-paper or online settings (*16*). It has also previously been used in evaluations of SMART (*12*). The summative score is the total of all five items (5–35) with higher scores indicating higher life satisfaction. This score can be broken down categorically into Extremely Dissatisfied (5–9), Dissatisfied (10–14), Slightly Dissatisfied (15–19), Neutral (20), Slightly Satisfied (21–25), Satisfied (26–30), and Extremely Satisfied (31–35).

Relevant narrative comments described in the above Education Outcomes Evaluation were used to contextualize wellbeing questionnaire results. In particular, responses regarding Dialogue and SMART were reviewed, as these two sections were most intentionally focused upon wellbeing. As mentioned in the Discussion section of the main text, an external control group of previous Mayo Clinic summer undergraduate students was recruited to give further confidence to study wellbeing findings. The education measures were not administered to this external control group given that they had already attended Mayo Clinic's summer undergraduate research program in 2019.

<u>Delivery</u>

**IRB**.

The study including wellbeing surveys and contact materials was reviewed and approved by the institution's COVID-19 committee and Education committee. Subsequently, an Institutional Review Board member reviewed and determined the survey to qualify as minimal risk research exempt under 45 CFR 46.101 category 2. Assent was sought before the start of each wellbeing survey.

**Qualtrics**.

Surveys were delivered, administered, and responses recorded using Qualtrics software (Copyright 2020 Qualtrics, Provo, UT). Surveys were delivered on the first and last days of the program and remained open for one week to maximize response rate. The education surveys and wellbeing surveys were delivered and completed independently. Remuneration was granted for the completion of the wellbeing surveys, though participants were free to leave questions blank. The education survey was considered a required course component.

##### Quantitative and Qualitative Analysis

<u>Quantitative Analysis</u>

**Education:** As a result of embedding education outcomes questionnaires into the main program and accompanying stipend, 100% of the participants who engaged in and completed all required parts of SFIR responded to the surveys. For items related to confidence in research skill and career knowledge and interest, trends were analyzed by comparing distribution of responses from pre- to post- program. Items were grouped together for analysis based on the original inventory's categorization, resulting in five dimensions: conceptualizing a study (3 questions), designing a study (4 questions), collaborating with others (2 questions), protecting research subjects and responsible conduct of research (2 questions), and design visual presentations (1 question). A small number of students left empty responses that precluded summative scores. Data presented in Figure 1 are unmatched and include all students with pre- or post- responses. Items requesting participant feedback about the various program components were analyzed by looking at the distribution of Likert scale ratings of those components.

**Wellbeing:** Wellbeing survey opt-in for SFIR students was 81%. Analysis was performed in JMP Pro 14 and only included respondents who attended the entirety of the SIFR program and who provided complete pre- and post- surveys for each given inventory. Conservative non-parametric tests (Wilcoxon Signed Rank) were run on the summative scores and compared to their parametric counterparts (paired t-test). As statistical significance did not noticeably change and responses displayed adequate normality in addition to sufficient sample size, paired t-test values were reported for ease of comparison. Effect size calculations are paired samples t-test Cohen's d, derived by dividing the mean difference by the standard deviation of the paired difference, $d = \frac{mean_D}{SD_D}$. Interpretation follows standard guidelines set by Cohen, where d = 0.2, 0.5, and 0.8 correspond to small, medium, and large effect sizes, respectively (*17*). For a more practical view of results, wellbeing responses were binned utilizing established inventory cut-offs above and visualized as a distribution for trend analysis. Survey opt-in for SURF controls was 63% (N=36). Analysis was performed in Prism 8.43 and only included respondents who provided complete pre- and post- surveys for each given inventory. Due to relatively small external control sample size and potential skewness in distribution, non-parametric tests (Mann-Whitney U) were run on the summative scores.

<u>Qualitative Analysis</u>

Two independent investigators (KK and AMG) separately conducted typological analysis on narrative data in the eight weeks following the program's conclusion. The data were binned into two typologies: (1) positive or complimentary feedback and (2) critical or mixed feedback. Entries were re-read by typology, and main ideas were summarized. Patterns within each typology were identified and recorded, and the data was re-examined for non-examples. Non-examples found were largely related to suggestions for alterations to the delivery of the program component under study. Patterns were summarized as one-sentence generalizations and compared between the two investigators, as well as with a third senior investigator (LMS). Representative data excerpts supporting the generalizations were identified and agreed upon by the investigators and included in the study results. See Supplementary Material 6 for more details.

#### REFERENCES


1. C. Pierret *et al.*, Improvement in Student Science Proficiency Through InSciEd Out. *Zebrafish* **9**, 155-168 (2012).
2. E. A. Mullikin, L. L. Bakken, N. E. Betz, Assessing Research Self-Efficacy in Physician-Scientists: The Clinical Research APPraisal Inventory. *Journal of Career Assessment* **15**, 367-387 (2007).
3. G. F. Robinson *et al.*, A shortened version of the Clinical Research Appraisal Inventory: CRAI-12. *Acad Med* **88**, 1340-1345 (2013).
4. B. W. Smith *et al.*, The brief resilience scale: assessing the ability to bounce back. *International journal of behavioral medicine* **15**, 194-200 (2008).
5. E. M. E. Smith B.W., J. Alexis Ortiz, Paulette J. Christopher, Erin M.Tooley., in *Resilience in Children, Adolescents, and Adults,* S. D. Prince-Embury S., Ed. (Springer,, New York, NY., 2013).
6. G. Windle, K. M. Bennett, J. Noyes, A methodological review of resilience measurement scales. *Health and Quality of Life Outcomes* **9**, 8 (2011).
7. A. Bhagra *et al.*, Stress Management and Resilience Intervention in a Women's Heart Clinic: A Pilot Study. *Journal of Womens Health* **28**, 1705-1710 (2020).
8. S. Cohen, in *The social psychology of health.* (Sage Publications, Inc, Thousand Oaks, CA, US, 1988), pp. 31-67.
9. S. Cohen, T. Kamarck, R. Mermelstein, A Global Measure of Perceived Stress. *Journal of Health and Social Behavior* **24**, 385-396 (1983).
10. K. T. Cohen S, Mermelstein R, et al., *Measuring stress: A guide for health and social scientists*. S. Cohen, R. C. Kessler, L. U. Gordon, Eds., Measuring stress: A guide for health and social scientists. (Oxford University Press, New York, NY, US, 1997), pp. xii, 236-xii, 236.
11. E. H. Lee, Review of the psychometric evidence of the perceived stress scale. *Asian Nurs Res (Korean Soc Nurs Sci)* **6**, 121-127 (2012).
12. S. S. Chesak *et al.*, Stress Management and Resiliency Training for public school teachers and staff: A novel intervention to enhance resilience and positively impact student interactions. *Complementary Therapies in Clinical Practice* **37**, 32-38 (2019).
13. L. N. Dyrbye *et al.*, The Impact of a Required Longitudinal Stress Management and Resilience Training Course for First-Year Medical Students. *Journal of General Internal Medicine* **32**, 1309-1314 (2017).
14. *Assessing well-being: The collected works of Ed Diener*. E. Diener, Ed., Assessing well-being: The collected works of Ed Diener. (Springer Science + Business Media, New York, NY, US, 2009), pp. xi, 274-xi, 274.
15. E. Diener, R. A. Emmons, R. J. Larsen, S. Griffin, The Satisfaction With Life Scale. *Journal of personality assessment* **49**, 71-75 (1985).
16. R. T. Howell, K. S. Rodzon, M. Kurai, A. H. Sanchez, A validation of well-being and happiness surveys for administration via the Internet. *Behavior Research Methods* **42**, 775-784 (2010).
17. J. Cohen, *Statistical Power Analysis for the Behavioral Sciences*. (Academic Press, 1988), pp. 490.

Supplementary Table 1 – External Control Summary

| | | N | Pre (x̃ (IQR)) | Post (x̃ (IQR)) | Δ (x̃ (IQR)) | p-value |
|---|---|---|---|---|---|---|
| BRS | SFIR | 129 | 3.33 (2.83–3.83) | 3.5 (3.17–4) | 0.17 (-0.17–0.5) | 0.03* |
| | Control | 33 | 3.5 (2.83–4) | 3.5 (2.75–4) | 0 (-0.17–0.17) | |
| PSS | SFIR | 125 | 21 (15–25) | 18 (13.5–22) | -1 (-5–1) | 0.03* |
| | Control | 32 | 18.5 (13.5–22) | 19 (12.25–23.75) | 0 (-0.75–1) | |
| SWLS | SFIR | 131 | 25 (20–29) | 26 (21–30) | 1 (-1–4) | 0.81 |
| | Control | 32 | 24 (19.25–28) | 27.5 (20–29.75) | 1 (-1–4) | |

Table legend: Wellbeing response comparison of SFIR students and external control. Median and IQR are reported with Mann Whitney U test p-values comparing change. SFIR students' pre-post gains in resilience and stress remain statistically significant in comparison to the control cohort.

**Demographic summary of enrolled participants**

| | Gender | |
|---|---|---|
| | **Woman** | **Man** |
| **Percent (count)**<br><br>Note: none responded "Other" or "Prefer not to respond." | 37% (62)<br><br>FYI 48 SURF +14 UREP | 63% (106)<br><br>FYI 86 SURF +20 UREP |
| | **Historically Underrepresented in Biomedical Science** | |
| | **Yes** | **No** |
| **Percent (count)**<br><br>Note: n=1 unknown was counted as "No." | 28% (47)<br><br>FYI 46+1 | 72% (121)<br><br>FYI 88+33 |
| | **Disability status** | |
| | **Yes** | **No** |
| **Percent (count)** | 4% (7)<br><br>FYI 7+0 | 96% (161)<br><br>FYI 127+34 |

Demographics are not available for n=2 students not recruited through SURF or UREP and are excluded from this summary. For all reported variables n=168.

### SFIR Dialogue #1 Template

#### ***“I Hear You”***

*Covid and Me: In this Dialogue (only 20 min), we will focus closely on just individual experiences to the Covid shutdowns. We will steer out into our community during our next talk.*

Increasing pace and risk

Progress toward exit question

Increasing pace and risk

#### Entry Questions

- In what ways has your life looked different in the past months than it did a year ago?
- What changes have you made in your daily routine?

#### Transition Questions

- What activities are you part of now that include other people?
- What format is used for those interactions (face to face, digital, etc.)?
- Do you dress differently than you did before March?
- How has your diet changed in the Covid environment?
- How about haircuts- during the shutdown what was your solution? Did you keep the changes once the salons opened up?
- Have you experienced financial change during Covid?
- How would you say your personal levels of stress and anxiety have played out during the Covid experience?
- How has your health been?

#### Exit Questions

- As you imagine yourself in a year, how will you describe yourself during Covid?

**SFIR Dialogue #2 Template**

***"I Hear You"***

*COVID and My Science: In this discussion we will talk about the ideas in science that interest you. What movies are you seeing? What blogs do you follow? Has COVID changed your media consumption?*

Increasing pace and risk

Progress toward exit question

#### Entry Questions

- What was the last "sciency" movie you saw?
- What movies or books have you pulled back off the shelf during sheltering in place? Any of them pandemic related?

#### Transition Questions

- We are grouped today as a "track", but that is not all that would define your science. What ideas in science are really interesting to you right now?
- Why did you get interested in those areas?
- Do you have a "face to your science"? - Meaning do you think of yourself or someone you know when you think of the science that interests you?
- Who do you picture when you think of COVID19?
- Have you thought about how your science interests relate to COVID? Tell me about it.
- What would you want to do to add to what is known about coronaviruses?
- What kind of link is ok between funding of science and current events like the pandemic?
- Is there a danger in connecting funding of science too closely to current events?

#### Exit Questions

- What influences do you think belong in the shaping of your science?

Increasing pace and risk

### SFIR Dialogue #3 Template
#### *"I Hear You"*

*COVID and My Community (COVID Karens): In this Dialogue, we will explore the community around us and how we, as a group have responded to the pandemic.*

Increasing pace and risk

Progress toward exit question

#### Entry Questions

- There have been a lot of stories about "COVID Karens", people who have been caught on video having a meltdown about having to wear a mask". What reactions have you seen in your community?
- Have you met anyone upset by masking?

#### Transition Questions

- What are your thoughts on masking in your community? Are there places that you think are more or less important for masking?
- What do you think is the source of argument about precautions for COVID?
- Who should decide what we do next?
- Who would you value joining us in the room to talk about preparing for COVID?
- What unique attributes of being you do you think relate to your response to the previous questions?
- What differences across people in your community may drive different responses?
- What would you like to see happen next to improve outcomes of the pandemic?
- What in your life would you be willing to give up to see that outcome happen?

#### Exit Questions

- What do you consider to be a reasonable loss from COVID 19? Think in terms of people, resources, opportunity,etc.
- What loss is unacceptable?

Increasing pace and risk

### SFIR Dialogue #4 Template

#### *"I Hear You"*

*My Science and My Community: Today we will come full circle in our Dialogues and link your scientific interests and your home community.*

Increasing pace and risk

Progress toward exit question

#### Entry Questions

- If I was to point to one of the presenters of this program and say "Wow, that's a great scientist," what attributes come to your mind?
- What have great scientists done?
- What, then is great science?

#### Transition Questions

- Who determines what science is done professionally?
- How does money play into deciding what science is done?
- How might politics play into deciding?
- What examples of that have you seen during COVID?
- Can your community help decide? How?
- What examples have you seen of the community exerting force on science in recent months?
- What benefits come from including the community in science?
- What challenges are there to including the community in science?
- Are there groups within your community that you believe are better tied to science and healthcare than others? Why?
- Is there a difference in expectation of connecting science to a community if you are a basic, translational, or clinical researcher?

#### Exit Questions

- How will you connect your community to the science you do now and in the future?

Increasing pace and risk

**Summer Foundations in Research (SFIR): 1-on-1 Mentorship Expectations**

Traditionally the SURF program is one of the most robust recruitment and publicity tools for the Mayo Clinic Graduate School of Biomedical Sciences and the Medical School. Beyond that is one of the **best opportunities currently available for Mayo Clinic graduate students, postdoctoral fellows, and junior faculty to gain valuable mentorship experience** as they build their academic careers. Though originally slated to be entirely canceled for the summer of 2020 due to the prevailing realities of the Covid-19 Pandemic, a new entirely online offering: Summer Foundations In Research (SFIR) has been created to fill these key voids for both the student interns and for the graduate school. While this program will certainly be very different from the normal SURF program **as all participants will be remote,** we endeavor to recreate, as possible, many of the same core experiences that are key to the in-person program. Most important of these is the ability to interact 1-on-1, or in small groups with scientific mentors like **you.** Below is a brief summary of the guidelines and what will be expected of you.

- From Thursday August 6th thru Friday August 14th (referenced as "Mentoring 1-7" in the attached syllabus) we have scheduled time for you to meet. **However, you are free to make whatever arrangements that work for you and your mentee anytime time during the program**. These mentoring sessions can be with a post doc, phd students or other faculty.
- We ask that you introduce them to and involve them in (as possible) your science. Invite them to participate in what you are currently doing; research, write, communicate, perform, or analyze a current experiment or project.
  - Examples of virtual activities may include:
    - Zoom meetings to discuss the science,
    - walking them through the literature search process,
    - showing them around your lab via video conference,
    - a video conferenced experiment,
    - involving them in the planning of an experiment,
    - having them help you analyze data, or anything you can think of…
  - These experiences can include capacity building in any area of your science from literature review to PCR
  - Assign a PhD Buddy to give a virtual walking tour of the lab and campus – Plummer Library, etc.
  - Develop a "**Meet the Professor**" sessions in which each faculty has an opportunity to share their research and any SFIR could attend. The sessions could be included in the shared calendar
  - Ask a student to share their 100-hour internship experience to explore other career options
  - Ask a postdoc to share their story and what they're doing now.
  - Dr. Scarisbrick is creating sessions on Tuesdays, 3:00-4:00 that any SFIR can attend:
    - 3-Minute Thesis
    - how to apply to grad school.
    - Face Transplant lecture by Dr. Samir Mardini
    - Faculty & student presentations for any student interested in Regenerative Science
  - During the Mentoring time the goal would be for them to learn enough about your science to feel comfortable giving a 5 minute talk on the project along with a simple poster for them to present at a virtual symposium at the conclusion of the program.

    - **The poster will include three sections- one on the didactic experience of SFIR, one on their small group design, and one on the science they learn with mentors.**
- Finally please invite your mentees to any virtual lab or track events that you will be attending to give them an experience of the culture of research at the clinic.

In addition to these opportunities with your students if you participate as a mentor in this program you will have the option of attending mentorship training classes with Dr. Bruce Horazdovsky to help you hone your mentoring abilities not only for this program but also for further mentorship in your career. Finally, attached to this note is a copy of the syllabus for the entire SFIR program so you can see what your mentees will be experiencing before and while they are working with you. You would be welcome to attend for any program components that interest you such as Dr. Sood's SMART Mindfulness program, or the community dialogues on the covid-19 pandemic. If any of these offerings interest you please email Zachary WareJoncas for the option to be enrolled.

**Supplement for qualitative analysis with examples (Mindfulness only)**

| Code | Description | Example |
|---|---|---|
| Positive | Any participant comments with exclusively complimentary features. | "I noticed a marked improvement in my own mental health and intend to employ as many of the tools as Dr. Sood gave us as possible in my daily life. This program component was invaluable." |
| (sub-theme) Practical utility | Participant comments that have to do with the current or expected use of SMART and/ or SMART skills in one's life. Three dimensions were identified that include *personal*, comments that have to do with expected use of SMART or benefits to oneself; and *professional*, which included comments on the value or use of SMART skills in science, research, and/ or future career. | "Mindfulness was incredibly useful because of how it gave me a different perspective on how to address stress and issues in my life."<br><br>"Dr. Sood's mindfulness sessions were a highlight of the program for me. He gave really concrete and valuable advice for improving relationship(s) and [how to] have a positive, well adjusted mindset. All of these are highly valuable in a scientific career." |
| (sub-theme) Praise | Includes participant comments that conveyed admiration for the *presenter*, appreciation for the *content*, or appreciation for the *relevance* of the material. | "I loved Dr. Sood and the Mindfulness content."<br><br>"This deserves its own category of worthwhile-ness. It was truly an invaluable experience that I would strongly recommend to continue doing" |
| Mixed or negative | Any participant comments with critical feedback; could include both complimentary and critical feedback ("Mixed"). Sub-themes included critiques of *content* and the *use of time*. | *"I didn't necessarily agree with the lectures, but I know it was worthwhile for students that needed it."*<br><br>*"I know that stress management is important;* |

| | | |
|---|---|---|
| | These represent relatively few comments (9). | *however, I would have rather used the time to connect with students and faculty at Mayo."* |
